# Do amplifiers of selection maximise average fitness?

**DOI:** 10.1101/2022.03.29.486264

**Authors:** Nikhil Sharma, Arne Traulsen

## Abstract

Evolutionary dynamics on graphs has remarkable features: For example, it has been shown that amplifiers of selection exist that – compared to an unstructured population – increase the fixation probability of advantageous mutations, while they decrease the fixation probability of disadvantageous mutations. So far, the theoretical literature has focused on the case of a single mutant entering a graph structured population, asking how the graph affects the probability that a mutant takes over a population and the time until this typically happens. For continuously evolving systems, the more relevant case is when mutants constantly arise in an evolving population. Typically, such mutations occur with a small probability during reproduction events. We thus focus on the low mutation rate limit. The probability distribution for the fitness in this process converges to a steady-state at long times. Intuitively, amplifiers of selection are expected to increase the population’s mean fitness in the steady-state. Similarly, suppressors of selection are expected to decrease the population’s mean fitness in the steady-state. However, we show that another category of graphs, called suppressor of fixation, can attain the highest population mean fitness. The key reason behind this is their ability to efficiently reject deleterious mutants. This illustrates the importance of the deleterious mutant regime for the long-term evolutionary dynamics, something that seems to have been overlooked in the literature so far.

## I. INTRODUCTION

Understanding how spatial structures can affect evolutionary dynamics has been of interest to evolutionary biologists for a long time. More than a decade ago, a framework known as Evolutionary graph theory has been introduced [1]. The primary quantity of interest has been the fixation probability of a mutant on graphs, which is the probability that a mutant with given fitness takes over the rest of the wild-type population [2–6].

Fixation probability is a central concept in evolutionary biology, as it determines the rate of evolution in the low mutation rate regime [7, 8]. Spatial structure tweaks the strength of selection and genetic drift [9, 10]. As a consequence, some graphs have higher probability of fixation than others for a mutant with a given fitness value. Of particular interest are those graphs that increase the fixation probabilities for advantageous mutants and decrease the fixation probabilities for disadvantageous mutants – so called amplifiers of selection. While initially amplifiers seemed to be special structures [1], it turned out that under Birth-death updating, most random networks are amplifiers of selection [11].

However, fixation describes evolutionary dynamics only on a relatively short time scale. In the long run, additional mutants will arise and the population will eventually reach a steady state in terms of fitness. A model that investigates such long evolutionary trajectories in graph structured populations has been missing so far. This problem has been studied for two types [12], but not for the case of continuously arising mutations with a potentially infinite number of types. The neutrality counterpart of this problem has been investigated in [13] where equilibrium properties are shown to be independent of the population structure.

In Evolutionary graph theory, every node of a graph is an individual. If an individual produces an offspring, the offspring is placed in a neighboring node. We focus here on graphs that are undirected and unweighted, such that all neighboring nodes are chosen with the same probability. Graphs can be classified based on their fixation probability as compared to that of the complete graph. The fixation probability of a mutant with fitness *f*′ appearing in a population with fitness *f* on a complete network *C* (corresponding to a well-mixed population) is given by

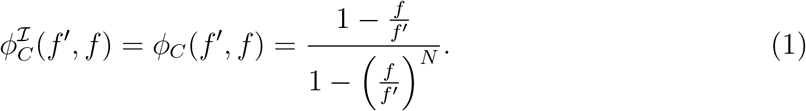

In this case, all nodes are equivalent and it does not matter where the mutation occurs. In the general case, the fixation probability depends crucially upon the node where the mutant first arises [14]. In Eq. 1, 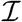 denotes this initialisation scheme, according to which a mutant arises on the network. To be specific, 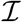 is the probability distribution represented by a row vector of size *N* where element *i* is the probability with which a mutant arises at node *i*. In the simplest case, this is a uniform probability – this case is typically referred to as uniform initialisation.

Based on fixation probabilities and the initialisation scheme, graphs typically fall into three categories [11]: Amplifiers of selection, suppressors of selection, and isothermal graphs:

- An *Amplifier of selection* is a graph *G* that increases the fixation probability of an advantageous mutant and decreases the fixation probability of a disadvantageous mutant compared to a complete graph, 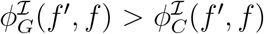 for *f*′ > *f* and 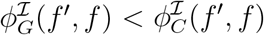 for *f*′ < *f*.
- A *Suppressor of selection* is a graph *G* that decreases the fixation probability of an advantageous mutant and increases the fixation probability of a disadvantageous mutant compared to a complete graph, 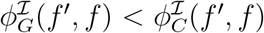 for *f*′ > *f* and 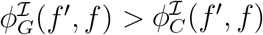 for *f*′ < *f*.
- An *Isothermal graph* is a graph G that has the same fixation probability as the complete graph, 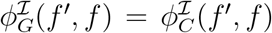 for all *f*′ and *f*. In particular, any graph where the number of links for each node is the same is an isothermal graph for uniform initialisation (in the more general case of weighted graphs, where mutants are placed on neighbouring nodes with different probabilities [15], the temperature of a node *i* is equal to the sum of the incoming link weights, 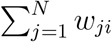).

For later on purposes, we introduce a fourth type [11, 16], Suppressors of fixation:

- A *Suppressor of fixation* is a graph *G* that reduces the fixation probability for both advantageous and disadvantageous mutants, 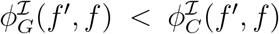 for all *f*′ ≠ *f*. These graphs are also called reducers of fixation [17].

The definition for isothermal networks has originally been defined for a uniform initialisation scheme. In general, such a definition would depend on the details of the update rule [18]. The classifications for amplifiers and suppressors above have also been developed for uniform mutant initialisation. For other initialisation schemes, for example, temperature initialisation, the classifications become less straightforward, especially near neutrality [19].

## II. MODEL

In order to generate any evolutionary dynamics, we must choose an update mechanism. We focus on the Birth-death (Bd) update rule, where first an individual is selected at random, but proportional to fitness. This individual produces an offspring, which is placed in one of the neighbouring nodes. We assume that the offspring is mutated with a small probability *μ* and identical to its parent with probability 1 – *μ*. The state of a population is represented by a fitness vector **f** = (*f*_1_, *f*_2_, ⋯, *f*_*N*–1_, *f_N_*)^T^, where *f_i_* is the fitness of an individual at node *i*. When an offspring is mutated, we choose its fitness *f*′ from a continuous bounded distribution *ρ*(*f*′, *f*), where *f* is the parent’s fitness and *f*_min_ ≤ *f*′ ≤ *f*_max_, see Fig. 1 (a).

**FIG. 1.**
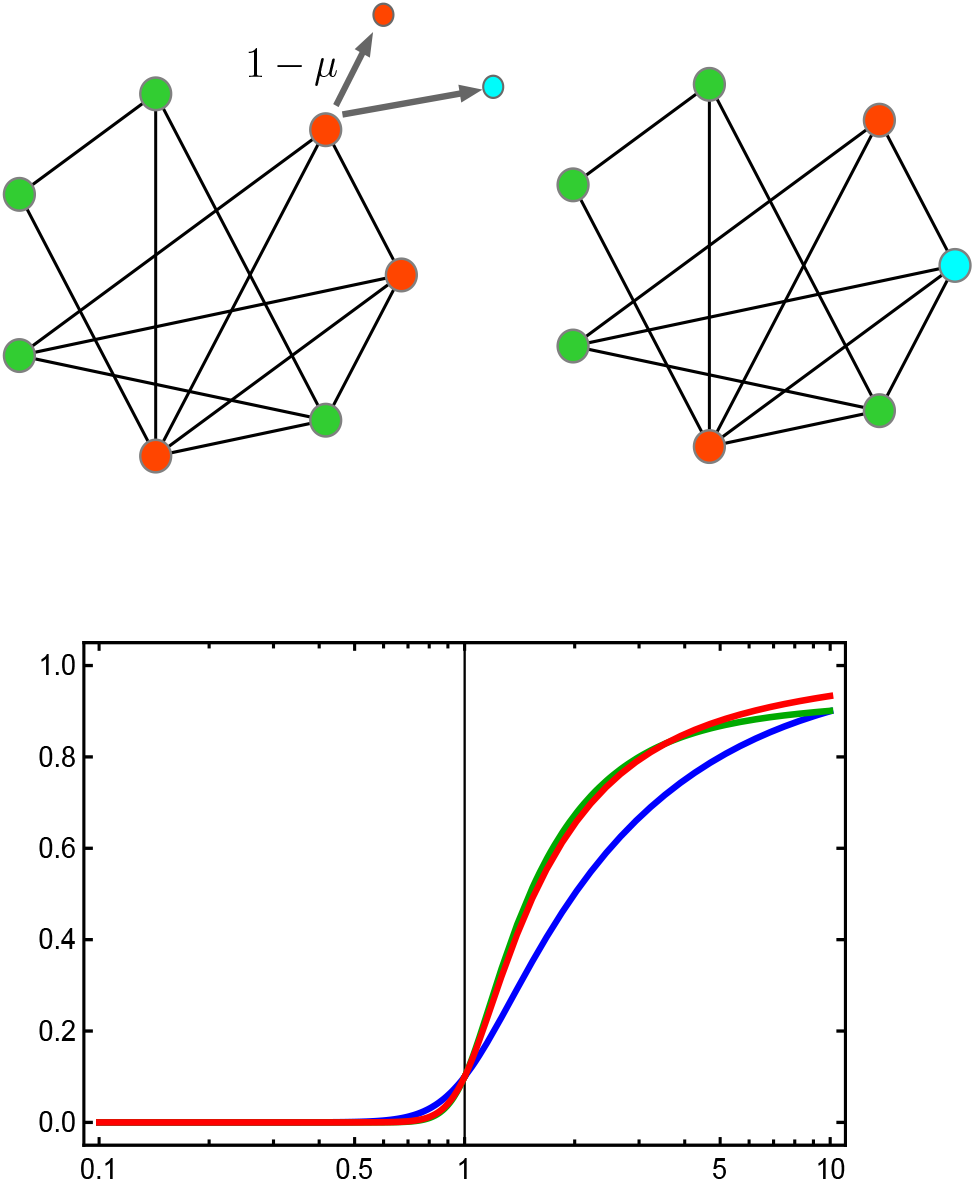
Moran Birth-death (Bd) with continuous mutation on a graph. (a) The Moran Birth-death (Bd) update mechanism with continuous mutation is shown for a small graph. An individual is selected to reproduce with probability proportional to its fitness. The offspring mutates with probability *μ*, and its fitness *f*′ is sampled from a distribution *ρ*(*f*′, *f*) where *f* is the parent’s fitness. A neighboring individual is then chosen for death at random among the neighboring nodes and the offspring is placed in the empty node. We work in the low mutation rate approximation where *μ* is very small, such that only a single type is typically present in the population. (b) Evolutionary dynamics in fitness space. For low mutation rates, the evolutionary dynamics effectively becomes a biased random walk on the fitness space. The transition rates depend on the mutation rate *μ*, the fitness distribution of the offspring *ρ*(*f*′, *f*) (shown in grey), and the fixation probabilities 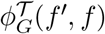. (c) The fixation probabilities 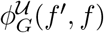 as function of mutant’s fitness *f*′ with *f* = 1 for uniform initialisation for three graphs with *N* = 10: complete, star and (weighted) star with self-loops. For this initialisation scheme, both the star and the star with loops are amplifiers of selection and all fixation probabilities intersect at *f*′ = *f* and 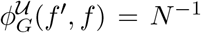. (d) Fixation probabilities as panel (c), but for temperature initialisation. Now the star with self-loops is only a piecewise amplifier of selection, as it reduces the fixation probability for very large *f*′. The star is a suppressor of fixation and reduces the fixation probability for all mutant fitness values. Note that even under neutrality, *f*′ = 1, the fixation probabilities are different.

Instead of looking into the full evolutionary trajectory of the system, we focus only on the fitness distribution in the steady state. As this is difficult for arbitrary mutation rates, we concentrate on the low mutation rate regime here. In this regime, the entire population effectively moves as a point (most of the time) in the fitness space. Typically, every individual in a population has the same fitness. Thus, the state of the population can be labeled by a single fitness value *f*. This is a good approximation if the time between two successive mutations is sufficiently high that a new mutant either gets extinct or takes over the entire population before the next mutation arises. Thus, our model describes sequential fixation [20]. We are mainly interested in the dynamics of probability density *P_G_*(*f,t*). One important observation is that mutations do not arise in all nodes with uniform probability, but mostly in those nodes that have more incoming links. This is captured by the idea of “temperature initialisation” [19], where the probability that a mutation arises is proportional to the temperature of a node *i*, in the case of unweighted networks 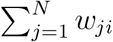, where *w_ji_* = 1 if a link between *j* and *i* exists.

With this, we can write down the master equation of the corresponding jump process [21, 22],

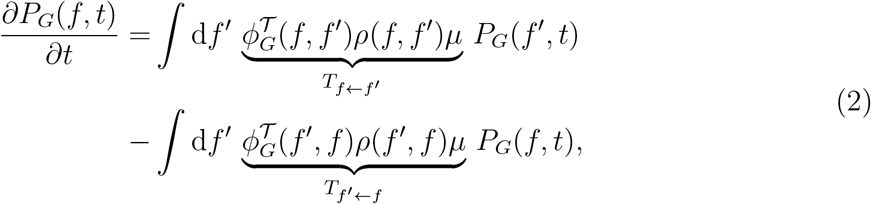

where *T_f′←f_* is the probability to change the population’s fitness from *f* to *f*′. *T_f′←f_* is given by the product of the probability of mutation, *μ*, the probability that a mutant with fitness *f*′ arises, *ρ*(*f*′, *f*), and the probability that such a mutation, arising preferentially in nodes that are replaced often, reaches fixation, 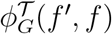 (where 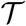 denotes temperature initalisation) The probability *T_f←f′_* is given by an equivalent argument.

The reason behind using fixation probability under temperature initialisation is that the probability for a mutant to arise on a certain node during reproduction in a homogenous population is proportional to the temperature of that node [19]. The corresponding evolutionary dynamics on the fitness space is a biased random walk with transition rates *T_f′←f_* and *T_f←f′_*. To derive the steady state, we start from the assumption of detailed balance [22],

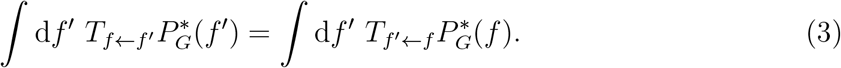

This condition is fulfilled if we have 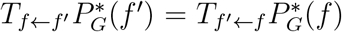 for all *f*, *f*′. From this, we obtain the steady state [22],

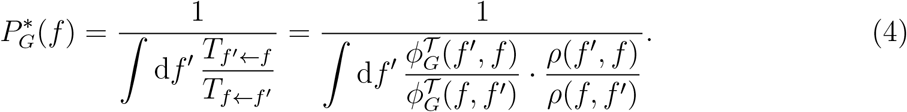

If the mutant’s fitness distribution *ρ*(*f*′, *f*) only depends on the fitness distance between parent and offspring, *ρ*(*f*′, *f*) = *ρ*(|*f*′ – *f*|), the population’s steady-state statistics depends only on the fixation probabilities - in this case, the second factor cancels out. Furthermore, the steady-state probability density for the population’s fitness is the same for all isothermal graphs, irrespective of the offspring mutational fitness distribution - the first factor is the same for all isothermal graphs.

To study mutation-selection balance on graphs, we choose the complete graph, the star, and the star with self-loops, i.e. a star where every individual can also be replaced by its own offspring. For finite *N*, under temperature initialisation a complete graph is an isothermal graph, while the star is a suppressor of fixation and star with self-loops is a piece-wise amplifier of selection [19] (and only for *N* → ∞, it becomes an amplifier in a strict sense), see Fig. 1 (c),(d). Our reason for using these graphs is because exact expressions for the corresponding fixation probabilities are known (see App. VB). The fixation probabilities for these three graphs are plotted as a function of mutant’s fitness in Fig. 1 (d).

The microscopic Moran-Bd update is costly to simulate for low mutation rates, especially for large graphs. Thus, we use a coarse grained description where we focus on the changes arising from mutations. We use the following Monte-Carlo type algorithm, which measures time steps in terms of mutational events,

- An initial fitness value *f* between *f*_min_ and *f*_max_ is assigned to each individual in a population on a graph *G*.
- At every time step, a mutant *f*′ is drawn from a distribution *ρ*(*f*′, *f*).
- The fitness of the entire population is then updated with probability 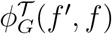 and with probability 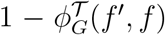 it remains the same. Note that this step takes into account that mutants tend to arise in different places with different probabilities.
- The last two steps are repeated for a sufficiently long number of time steps until a mutation-selection balance is attained.

We use this algorithm to infer how mutation-selection balance is attained in our three spatial structures. In Fig. 2 (a), the average fitness of the population is plotted as a function of the number of mutations occurred.

**FIG. 2.**
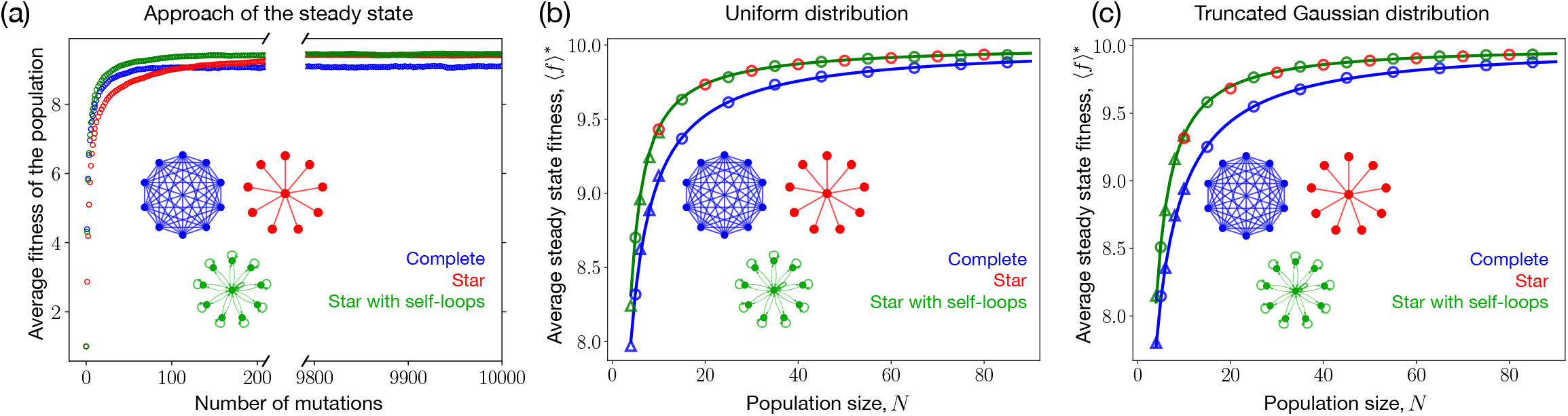
Average fitness trajectories and steady state fitness for different graphs. (a) The average fitness trajectories for the complete graph, the star graph, and the self-looped star graph with a uniform mutational fitness distribution, starting from a population where every individual has fitness *f* = 1. Although it is a suppressor of fixation, the star graph attains the same steady state fitness in the mutation-selection balance as the self-looped star graph. This is due to its better response to deleterious mutants (see inset of Fig. 1 (d)). (b) and (c) Steady state fitnesses in the mutation-selection balance attained for the self-looped star graph (green) and complete graph (blue) as a function of population size are shown in with (b) uniform and (c) truncated Gaussian mutational fitness jump distribution. The standard deviation for the Gaussian distribution is chosen to be *σ* = 1. Solid lines are the numerical solutions of Eq. 4, circles represent simulation points obtained using the Monte-Carlo algorithm, while triangles correspond to microscopic Moran Bd simulations, which are feasible only for very small *N*. Regardless of the mutational fitness jump distribution, the self-looped star (an amplifier) always attains higher steady state fitness in the mutation-selection balance than isothermal graphs for all (finite) *N*. Red circles correspond to the steady state fitness attained by the star graph, a suppressor of fixation. From (b) and (c), we find that even for large *N*, the star graph reaches almost the same steady state fitness as that of the self-looped star graph (parameters *N* = 10, *f*_min_ = 0.1 and *f*_max_ = 10, averages over 2000 realisations).

## III. RESULTS

The star with self-loops, an amplifier of selection, reaches the highest steady state fitness in its mutation-selection balance, see Fig.2 (a). This is expected because, compared to the other graphs, the self-looped star graph is best at fixing beneficial mutants for temperature initialisation. What is surprising is that the star graph, a suppressor of fixation with lower fixation probability for all fitness values of a mutant, not just attains higher steady state fitness than the complete graph (and thus all isothermal graphs), but almost the same balance as the star with loops. We also observe that just like isothermal networks, the star graph with loops takes fewer mutations to reach the balance than the star graphs, which takes many more mutations to reach the steady-state. In the subsequent sections, we will discuss these observations and give an explanation for them.

### A. Amplifiers attain higher steady state fitness in mutation-selection equilibrium

Amplifiers of selection are better at fixing beneficial mutants and avoiding the fixation of deleterious mutants than isothermal graphs. They are expected to attain a higher steady state fitness in the mutation-selection balance than isothermal graphs. We verify this expectation in Fig. 2 (b) and (c), which shows the average steady-state fitness of the population for different population sizes. We consider two types of mutational fitness jump distribution, a uniform and a (truncated) Gaussian, centered around the parent’s phenotype. We observe that regardless of the mutational fitness distribution, the star graph with loops attains a higher steady state fitness than the complete graph for all considered population sizes. In App. V A, we provide formal a proof that for any mutational fitness distributions, amplifiers of selection have higher steady-state fitness than well-mixed populations and suppressors of selection.

In the limit of very large graphs, *N* → ∞, the steady state fitness approaches the maximal possible value, 〈*f*\*〉 → *f*_max_, for both complete as well as self-looped star graph. In fact, *f*_max_ becomes an absorbing state for the evolutionary trajectories on these graphs and thus, the fluctuations around the steady state also goes to 0. This happens because for very large *N*, the fixation probability for the self-looped star under temperature initialisation becomes 1 – *f*^2^/*f*^′2^ for any *f*′ > *f* and 0 otherwise (see App. VB), whereas from Eq. 1 it follows that for complete graph it becomes, 1 – *f/f*′ for *f*′ > *f* and 0 otherwise. This also leads to the conclusion that the self-looped star needs fewer mutations than the complete graph to reach *f*_max_.

### B. Do amplifiers maximise average fitness?

The star graph with no self-loops, a suppressor of fixation, attains almost the same steady state fitness in mutation-selection balance as the star with self-loops, an amplifier of selection, as shown in Fig. 2 (b) and (c). Two main factors drive the steady state fitness: (i) fixing beneficial mutants with high probability and (ii) avoiding the fixation of deleterious mutants. As an amplifier of selection, the star with self-loops is superior in fixing beneficial mutants. But the star without self-loops is much better at avoiding the fixation of deleterious mutants, see inset of Fig. 1(d). In this way, the star without self-loops compensates for its lower fixation probabilities of beneficial mutants. However, this comes at a cost in terms of the time it takes to reach the steady state. There is additional perspective to this: Let us take two amplifiers *A*_1_ and *A*_2_, and without any loss of generality, assume that *A*_1_ is a better amplifier than *A*_2_. That is, *A*_1_ is better in fixing advantageous mutants and avoiding the fixation of deleterious mutants than *A*_2_. In that case, by the proof given in App. V A, it follows that the average steady state fitness of *A*_1_ is greater than the average steady state fitness of *A*_2_, i.e. 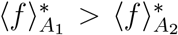. However, if *A*_2_ can be modified, along the lines of [23], such that its fixation probability for deleterious mutants becomes much lower than *A*_1_, the average steady state fitness 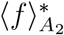 can exceed 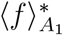.

### C. Time to reach the steady state

So far, we have studied the steady state average fitness values for different graphs. Now, we discuss the time they take to reach their respective steady states. To estimate these times, we use the concept of mixing times [24]. In a generic stochastic process, the probability distribution *P_G_*(*f*) defined on a space Ω changes towards its steady state. The mixing time can then be defined by tracking the distance of this evolving distribution to its steady state 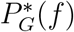 go to zero. Formally, the mixing time *t*_mix_ is defined as the minimal time when the distance *d*(*t*) to the steady state distribution is smaller than a threshold *ε*,

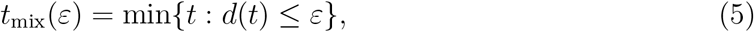

where,

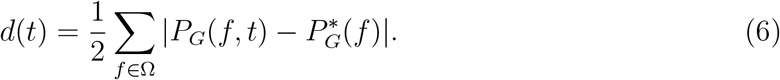

In Fig. 3 (a), we find that the star graph, a suppressor of fixation, attains the same balance as the self-looped star, but takes many more mutations to reach the steady-state than all other considered graphs. This is natural, as most mutations arising in this graph do not reach fixation. Another interesting observation is that for large *N*, the self-looped star graph has the smallest mixing time. This contrasts with the typical fixation probability and time relation, where larger fixation probability tends to correlate with higher fixation times [25–28]. Mixing time is difficult to calculate in the general case. But for single rooted graphs with with 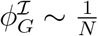, it can be computed efficiently. In App. V C, the mixing time is computed for this case.

**FIG. 3.**
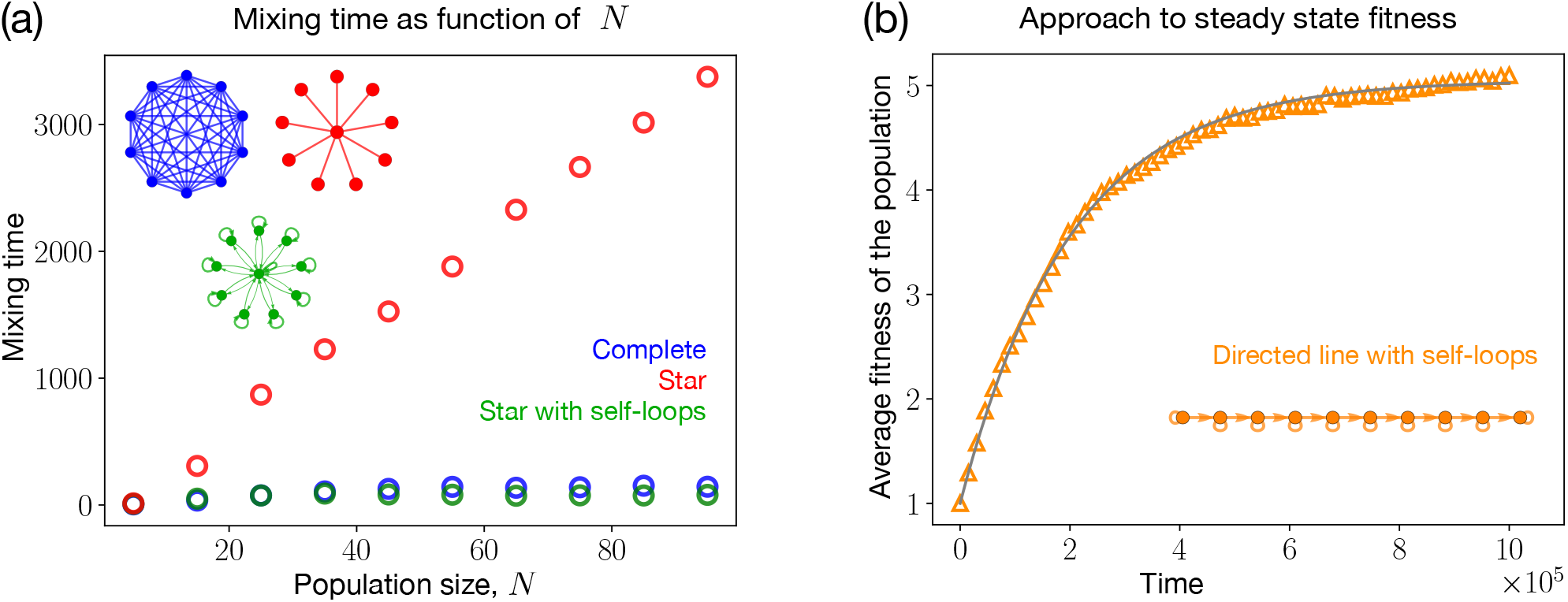
Approach to the steady state. (a) Mixing time for various graphs as a function of population size *N* measured in number of mutations. Parameters are as in Fig. 2. Mixing times have been obtained using Monte-Carlo algorithm. The star graph takes the longest to reach a steady-state. This is because the star graph, especially for large *N*, has very small fixation probabilities even for beneficial mutants and thus more attempts to increase fitness are needed to reach the steady state (For the mixing time, we assume 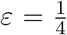 in Eq. 5). (b) Increase of average fitness on the directed line with self-loops. We compare the expression 27 and the corresponding microscopic Moran Bd simulations. Parameters are the same as in Fig. 2, with a mutation probability *μ* = 10^−4^. We see a good agreement between simulations (orange symbols) and the analytical result (gray line). From App. V C, we know that the relaxation time for single rooted graphs goes as 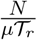. Substituting values for the parameters and the root temperature for the directed line with selfloops, 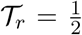, we get the mixing time to be approximately 2 × 10^5^. Here, the time is measured in terms of the number of Moran Bd steps. Although the simulation here is performed for the self-looped directed line, the same result is expected for any other single rooted (self-looped) graph as long as *μ* is sufficiently low. For more details, see App. V C.

### D. Gaussian fitness-phenotype map

Until now, the dynamics were considered in fitness space. However, in many cases mutations occur on the level of an individual’s phenotype. In that case, the state of a population is represented by a phenotype vector **p** = (*p*_1_, *p*_2_, ⋯, *p*_*N*−1_, *p_N_*)^T^, where *p_i_* is the phenotypic trait value of the individual at node *i*. The Moran Bd update still only takes the fitness of individuals into account. Therefore, a phenotype-fitness map *f*(*p*) is used to assign a fitness value to a phenotype. The fitness profile of the population then can be denoted by **f**(**p**) = (*f*(*p*_1_), *f*(*p*_2_), ⋯, *f*(*p_N_*))^T^. Similar to the previous case, when the offspring is mutated, we choose its phenotype *p*′ from a continuous bounded distribution *ρ*(*p*′, *p*), where *p* is the parent’s phenotypic value and *p*_min_ ≤ *p*′ ≤ *p*_max_. In this way, the work presented so far is identical what would obtain by considering a linear phenotype-fitness map, *f*(*p*) = *p*. In this section, we study the dynamics of a non-monotonic Gaussian map. In Fig. 4 (a), we show that the order of the steady state fitness in mutation-selection balance for different graphs remains the same as that of the linear phenotype-fitness map. It is interesting to note that the average steady state phenotype for all the graphs is the same, see Fig. 4 (b). Nevertheless, we see different steady state average fitness values. Even though all graphs have the same steady state phenotypic value, isothermal graphs are more prone to fluctuations and have a lower steady state average fitness value, see Fig. 4 (c). In a nutshell, this emerges because the average of a function in general is not equal to the function of average, i.e., 〈*f*(*p*)〉 ≠ *f*(〈*p*〉).

**FIG. 4.**
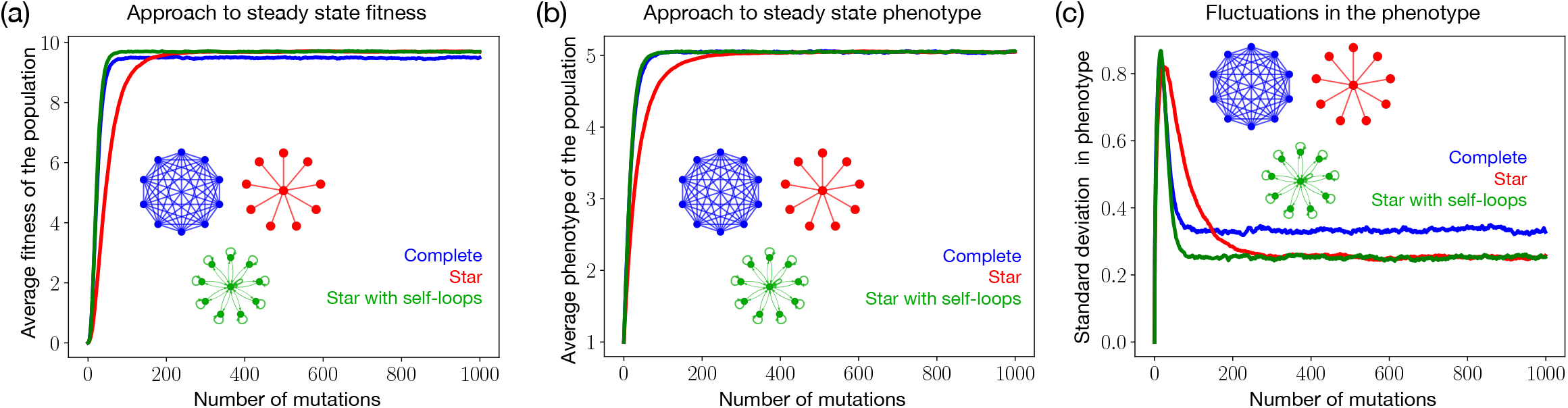
Gaussian phenotype-fitness map: Fitness and phenotypic trajectories for different graphs. Here, we consider a Gaussian phenotype-fitness map, 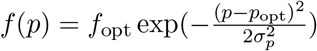, where *p*_opt_ is the optimal phenotype where the fitness is maximal and *f*(*p*_opt_) = *f*_opt_. The width of the map is *σ_p_*. (a) As in the case of a linear phenotype-fitness map, the star and the self-looped star have a higher state fitness than isothermal graphs. (b) All graphs converge to an average phenotype corresponding to the fitness maximum, but they have different fitness in steady-state (see panel a). The reason for this is are differences in the fluctuations in steady state. (c) The standard deviation of the phenotypes shows that the complete graph is more prone to fluctuations than the other two graphs (parameters *f*_opt_ = 10, *p*_opt_ = 5.05 and *σ_p_* = 1, all other parameters are as in the previous figures with *σ* = 0.5 for the Gaussian mutational jump phenotype distribution).

## IV. DISCUSSION

In Evolutionary graph theory, evolutionary dynamics on graph structures have been studied in great detail. While the field has been mostly driven from the mathematical and computational community [1, 2, 18, 27, 29], partly driven by biological systems [16, 30, 31], now there is increasing interest in applying these ideas to experimental systems in microbiology [32, 33].

The prospects of engineering a population structure that can optimise the chances to evolve certain mutations or to observe evolved population structures that minimise the evolution of mutations seem exciting, but these applications call for an extension of the field of Evolutionary graph theory: Most applications implicitly assume that each node is a small population and not all results carry over from graphs of individuals to graphs of sub-populations [34–37]. In addition, the field has focussed so far on fixation probability and fixation time [38–43].

This approach assumes that we can focus on the fate of a single mutant, but it can break down when mutations continuously arise, especially in graphs where the time to fixation or extinction is very high. Moreover, in the case where mutations continuously arise, one has to take into account where they arise. Thus, one needs to work with temperature initialisation, where the definitions of amplifiers and suppressors of selection seem to be less clear-cut.

We developed such a model that takes the continuous supply of mutations into account. We found that the prevention of deleterious mutants from fixing can be more important than increasing the chances of advantageous mutants in order to obtain a higher steady state fitness in the mutation-selection balance. In our case, the star, a suppressor of fixation for temperature initialisation, beats isothermal graphs and attains almost the same balance as a self-looped star, an amplifier. The cause for this is the ability of the star graph to prevent deleterious mutants much better from fixing than isothermal graphs. The deleterious mutants regime is usually overlooked in the literature while studying fixation probabilities by using large *N* arguments. However, here we have shown that the deleterious mutants regime is equally important, if not more, even for large *N*, as the beneficial mutants regime when studying long-term evolutionary dynamics.

Typically, amplifiers of selection have also a higher fixation time [27–29]. Thus, one has to be careful that the assumption of small mutation rate is still fulfilled. The weak mutation approximation works well if the time between two mutation events is much longer than the time to to reach fixation or extinction of mutant. Thus, the validity of this approximation depends not only on the mutation rates, but also the fixation times. Previous research has already indicated that that graphs with higher fixation times call for lower mutation rates for this approximation to hold [26, 28, 29]. The star graph with self-loops is very restrictive to the low mutation approximation compared to the complete graph, while for the star graph and the complete graph the weak mutation approximation starts to fail at similar mutation rates. This may be unexpected, as for uniform initialisation the fixation time in the star graph is an order of magnitude higher than in the complete graph. But here we observed that for temperature initialisation, the fixation time for the star graph becomes almost the same as that of the complete graph, while the fixation times for self-looped star remain much higher. However, if this assumption of weak mutation rates is fulfilled, structures that allow more mutations to reach fixation will have a smaller mixing time - therefore, amplifiers of selection tend to have a lower mixing time until their steady state fitness is reached.

Due to their ability to reach a high steady state fitness, population structures that suppress selection could be much more interesting in applications than previously thought — suppressing selection may be as relevant as amplifying it. We have shown that in a situation where a dynamic, graph-structured population continuously evolves, the amplification of selection via the promotion of advantageous mutations does not necessarily imply a higher steady state fitness. Instead, in such a process one has to carefully consider also the fate of deleterious mutations, an issue that has so far not been in the focus on Evolutionary graph theory.

## V. APPENDIX

### A. Amplifiers of selection attain higher steady state fitness in mutation-selection balance than suppressors of selection

Here we prove that amplifiers of selection attain higher steady state fitness in their mutation-selection balance than suppressors of selection. We denote an arbitrary amplifier by *A* and an arbitrary suppressor by *S*. From the definitions of amplifiers and suppressors mentioned in the introduction, we have for every *f*′ < *f* (and arbitrary initialisation scheme 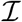)

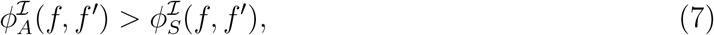

as well as

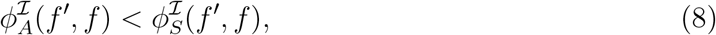

Combining these two inequalities, we get,

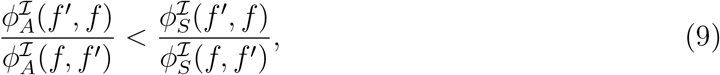

Multiplying the above equation with *ρ*(*f*′, *f*) /*ρ*(*f, f*′) followed by integrating over *f*′ from *f*_min_ to *f*, we obtain,

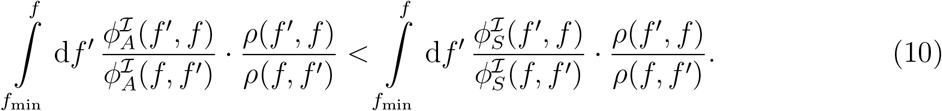

Similarly, we find,

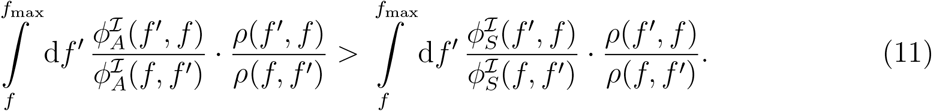

By making use of the steady-state solution Eq. 4 and Eq. 10, we find an inequality for probability density functions at the boundary *f*_max_ of the fitness domain,

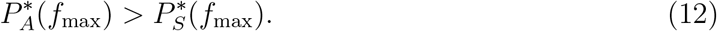

While Eq. 4 contains temperature initialised fixation probabilities, the same expression follows for the steady state statistics for any arbitrary initialisation. As an example, instead of the mutations taking place during reproduction they could appear spontaneously at any of the nodes. Eq. 4 would then contain uniform initialised fixation probabilities.

Similarly at the fitness boundary *f*_min_, using Eqs. 4 and 11 we have,

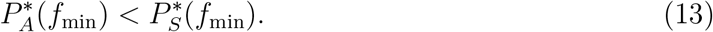

The very same inequalities, 12 and 13, hold if *A* (amplifier) or *S* (suppressor) is replaced by *C* (complete).

Let us now first prove that amplifiers of selection attain higher steady state fitness in mutation-selection balance than the complete graph. We take the case of an amplifier and the complete graph of the same size. The inequalities 12 and 13 imply that there exists a fitness, 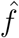, such that,

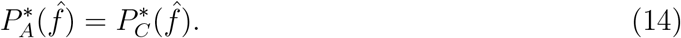

For simplicity here we assumed that there is only one intersection point for the curves 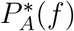 and 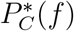. The proof, however, does not rely on this and can be extended to the general case where more than one intersection point are there. The main idea behind the proof has been sketched in Fig. 5. Let us define the 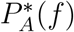 as:

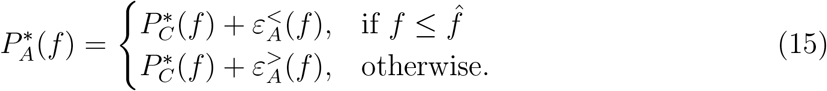

**FIG. 5.**
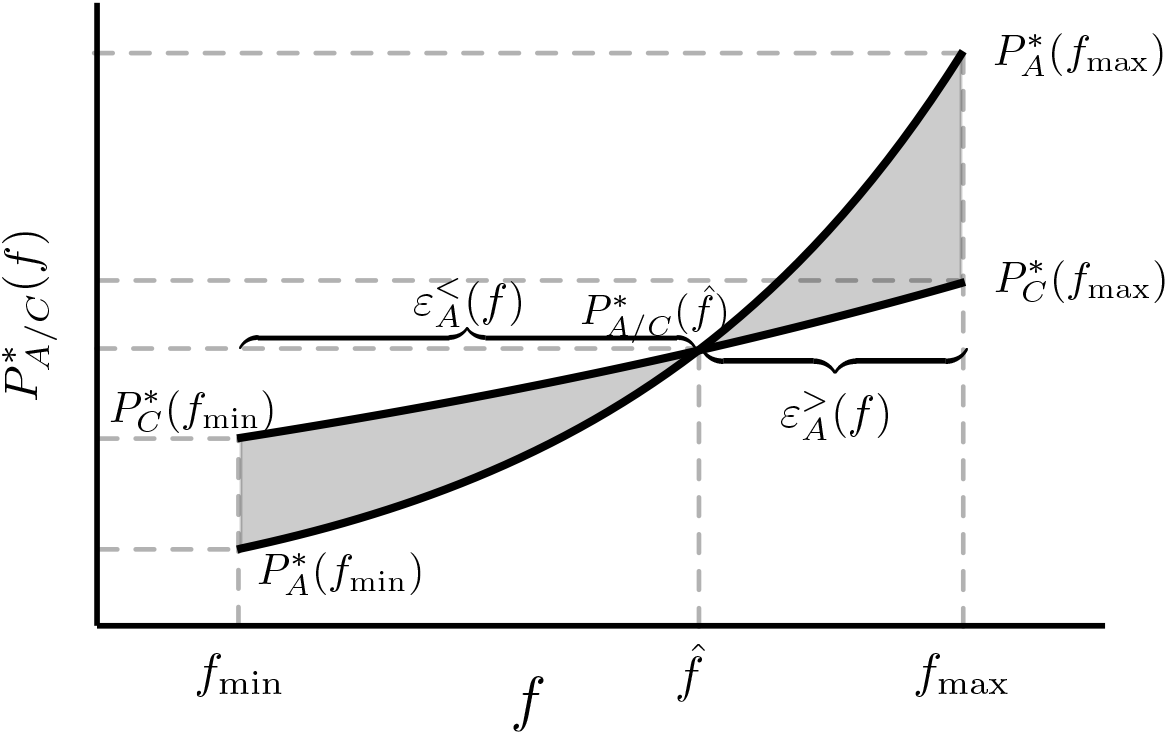
Sketch for the proof. Here we sketch the main idea behind the proof that the average fitness of an amplifier of selection exceeds that of a suppressor of selection or the complete graph in the steady state. Without any loss of generality, we take the case of an amplifier *A* and the complete graph *C*. The proof starts by first computing the order of 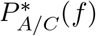 at the fitness boundaries *f*_min_ and *f*_max_. It turns out that 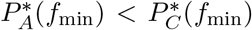 and 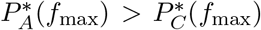. This implies that there exists a fitness point 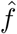 where these probability densities intersect. 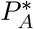 is then decomposed as the sum of 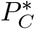 and the functions 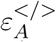, see Eq. 15. By using the normalisation conditions for 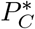 and the properties of functions 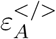, Eq. 16, we prove that amplifiers of selection attain higher steady state fitness than the well-mixed population and by extension, the suppressors of selection.

From the normalisation condition of 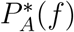 and 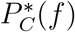, it follows that

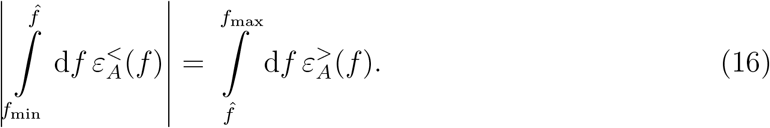

We are interested in the difference of mean fitness in the steady state,

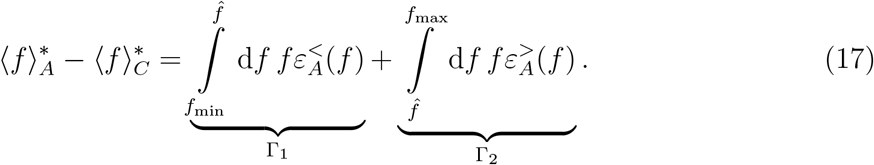

The first term in the above equation is negative, Γ_1_ < 0, while the second term is positive Γ_2_ > 0. In the following, we show that the magnitude of the term Γ_1_, is less than the term Γ_2_. Taking Γ_1_, we find that

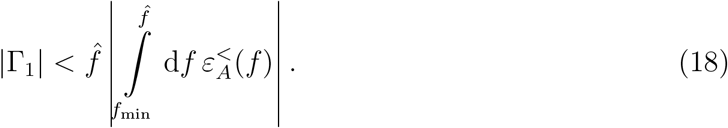

Similarly,

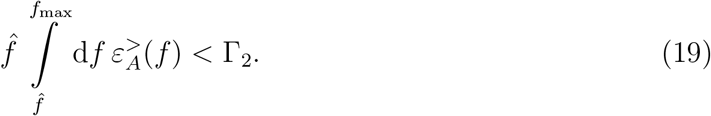

Now, because

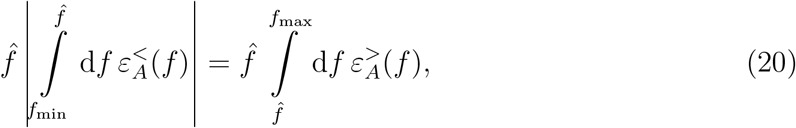

using inequalities 18 and 19, we find |Γ_1_| < Γ_2_. Therefore at last, we have

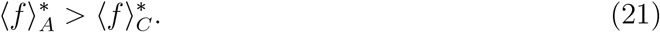

Following the same procedure, one can show that 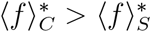. This implies that 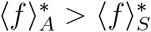 for all amplifiers of selection *A* and suppressors of selection *S*.

### B. Fixation probability for the self-looped star graph under temperature initialisation

In this section, we introduce the weighted self-looped star graph that has been used throughout the main text. It is defined by the weighted adjacency matrix

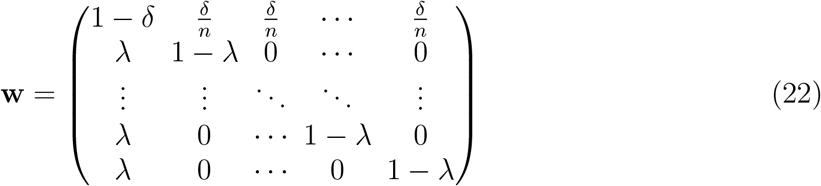

with 0 < λ ≤ 1 and 0 < *δ* ≤ 1. Here, *w_ij_* is the weight of the link directed from the node *i* to node *j* with the center being the node 0. The fixation probability under temperature initialisation for this graph has been derived by using the techniques of martingales [3, 19] and is given by

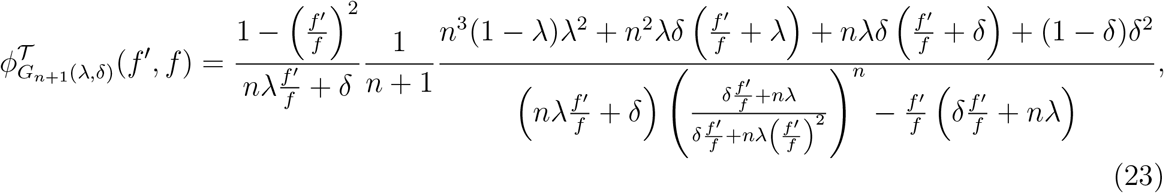

where *G*_*n*+1_(λ, *δ*) denotes the weighted self-looped star graph with *n* leaves. In the limit of *n* → ∞, when λ and *δ* are independent of *n*, Eq. 23 becomes,

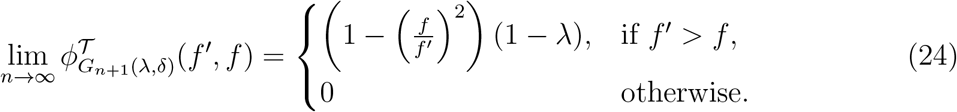

We use two versions of this weighted graph by first setting λ = *δ* = 1 in the above equation that results in the unweighted star graph which, however, is a suppressor in the limit *n* → ∞ under temperature initialisation. This is reflected by 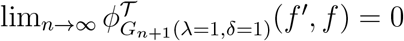 for all *f*′ and *f*. Setting instead 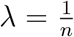 and 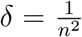 followed by taking the infinite population size limit of Eq. 23 yields a structure which is an amplifier for *n* → ∞. That is,

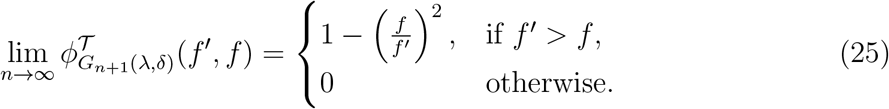

### C. Mixing time for single rooted graphs

For any one-rooted network with self-loops, we can use Eq. 2 to find an exact expression for the first moment when mutational fitness jump distribution is uniform. To do so, we first note that the fixation probabilities become independent of fitness values, 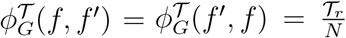, where 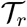 is the temperature of the root node. We then multiply Eq. 2 with *f* followed by integrating it over the entire fitness domain. Substituting the fixation probabilities, and using the normalisation condition for *P_G_*(*f*′, *t*), we obtain a first order ordinary differential equation for 〈*f*〉 in time

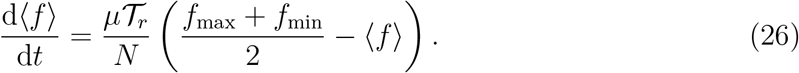

The average fitness for the case of any single-rooted graph (root with self-loops) with temperature 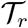 therefore changes in time as

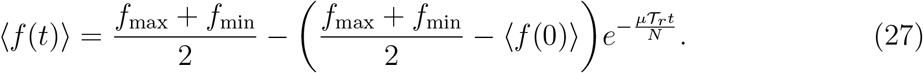

From this expression, we find that in the limit *t* → ∞, the fitness converges to the average of the fitness domain. Further, the mixing time, the number of mutations required to reach the steady-state, for these graphs scales as 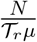. Therefore, the mixing time can be modulated by changing the temperature of the root node. We also note that the steady state for single rooted graphs (with self-loops) is independent of the choice of phenotype-fitness map — simply because the fixation probability for these graphs does not depend on fitness.

## ACKNOWLEDGMENTS

We are grateful to the members of the Evolutionary Theory department and the Dynamics of Social Behavior group for providing a very good scientific environment facilitating this work. NS sincerely thanks Carsten Fortmann-Grote for his assistance with computer cluster usage.

